# Drivers of Bird Communities in an Urban Neighborhood Vary by Scale

**DOI:** 10.1101/2024.01.21.576560

**Authors:** Andrea Darracq, Clay Bliznick, Ray Yeager, Jay Turner, Pradeep Prathiba, Jacob Pease, Howard Whiteman, Ted Smith, Aruni Bhatnagar

## Abstract

Given the accelerated pace of global biodiversity loss and rapid urbanization, it is becoming increasingly urgent to identify ways to minimize the costs and maximize the benefits of urban environments for wild flora and fauna. For instance, it has been estimated that 48% of all bird species are experiencing population declines. One of the main drivers of these declines is habitat loss and degradation associated with urbanization. Increased urbanization necessitates a better understanding of how to conserve birds in urban areas. Although relationships between urbanization and bird communities have been explored extensively, few studies have been conducted in residential neighborhoods, and the influence of urban environmental conditions, particularly air pollution, on bird communities remains unclear. In this study, we examined relationships between bird community metrics and environmental measures related to vegetation and air pollution within a residential neighborhood at multiple spatial scales. We found that bird species richness and the average number of native species were positively related to greenness (as measured by the normalized difference vegetation index; NDVI) within 50 m, and negatively associated with ambient levels of NO_2_ at 200 m. Similarly, we found the Hill-Shannon diversity index was positively associated with canopy cover, but negatively associated with NO_2_ at 200 m. The average number of invasive bird species, however, was negatively correlated with canopy cover at 50 m. The average number of native birds was negatively related to ultrafine particle (<100 nm in diameter) concentration. Unlike native bird abundances, invasive bird abundances were not sensitive to NO_2_ or ultrafine particles. Thus, our research suggests that reductions in air pollution, in combination with greening efforts that increase NDVI and canopy cover via the restoration of vegetation within urban neighborhoods, are likely to increase bird diversity and the abundances of native birds while reducing the abundance of invasive birds.

## Introduction

Recent assessments indicate more than 50% of the Earth’s land surface has been altered by humans (Hooke et al., 2012), leading to a profound loss of biodiversity and ecosystem collapse. These changes are occurring at an accelerated pace, and it has been estimated that current rates of extinction are 1,000 times greater than speciation rates (De Vos et al., 2015). Urbanization is one of the leading causes of land use alteration, and land use intensification associated with urbanization affects the spatial orientation of ecosystems, resulting in fragmented and degraded patches of urban green spaces with decreased connectivity, cover, and vertical stratification (F. Li et al., 2019; Z. Liu et al., 2016). Additionally, urbanization creates variation in air pollutants emitted by multiple sources (e.g., motor vehicles, power plants, residential heating, gasoline stations, consumer products; Karagulian et al., 2015; Liang & Gong, 2020; McDonald et al., 2018). Air pollution associated with urbanization and the resultant increases in greenhouse gases has not only contributed to changes in global climate (Romero-Lankao & Dodman, 2011; Rosenzweig et al., 2011), but it is also negatively affecting the health of living organisms, including humans (Kampa & Castanas, 2008; Newman, 1979; Singh et al., 2020; Work, 2022). Given the accelerated pace of global biodiversity loss and rapid urbanization, it is becoming increasingly urgent to identify ways to minimize the costs and maximize the benefits of urban environments for wild flora and fauna.

Previous studies have shown that wildlife (Aronson et al., 2014; Beninde et al., 2015; Turrini & Knop, 2015) and humans (Fong et al., 2018; Yeager et al., 2019) respond positively to greenness in urban environments. Increased greenness is associated with a more complex plant community relative to plant diversity and structure, which can support the needs of more wildlife species (Larue et al., 2018; Coverdale & Davies, 2023). Greenness can also facilitate greater abundances of primary consumers, such as arthropods, that may increase food availability for higher trophic levels (Fernández-Tizón et al., 2020; Lafage et al., 2014).

In addition to greenness, wildlife in urban environments are also affected by ambient air pollution. Common urban air pollutants include nitrogen dioxide (NO_2_), ozone (O_3_), carbon dioxide (CO_2_), sulfur dioxide (SO_2_), particulate matter (PM), and volatile organic compounds (VOCs) (Gulia et al., 2015). In humans, episodic exposure to air pollution has been shown to trigger acute events such as asthma attacks and myocardial infractions, where long-term exposures are associated with an increase in the risk of pulmonary and cardiovascular disease (West et al., 2016). Air pollution affects non-human animals as well. Several adverse health effects (e.g., respiratory distress or illness, increased glucocorticoids, behavioral immunosuppression, reduced reproductive success) have been observed in animals exposed to air pollution (Barton et al., 2023; Frutos et al., 2015; Newman, 1979; Sanderfoot & Holloway, 2017). Air pollution exposure has also been linked to reduced recruitment, survival, and diversity of animal populations (Barton et al., 2023; Newman, 1979; Sanderfoot & Holloway, 2017). For instance, increased levels of air pollution are associated with changes in the abundance and diversity of arthropod populations and reductions in arthropod sizes (Eeva et al., 1997; Pimentel, 1994; Zvereva & Kozlov, 2010), leading to reduced prey biomass.

Although landscape conversion, fragmentation, and exposure to air pollution are likely to influence most wildlife in urban environments, birds may be particularly sensitive. Approximately 20% of bird species occur in urban areas (Aronson et al., 2014). Moreover, bird communities are likely to be strongly affected by urban landscape features because nearly all birds are reliant on arthropods at some stage of their life cycle (Nyffeler et al., 2018) and bird species specialize in their use of the vertical strata of vegetative communities (e.g., understory, midstory, canopy; Finke & Snyder, 2008; Goetz et al., 2007; Huang et al., 2014; Jayson & Mathew, 2003; MacArthur & MacArthur, 1961), which are both influenced by the urban environment (Y. Liu et al., 2015; Meineke et al., 2023; Zhang et al., 2022). Birds may also be particularly vulnerable to air pollutants because of their respiratory system, which, unlike mammals, is characterized by unidirectional airflow and cross-current gas exchange (Bouverot, 1978; Brown et al., 1997). These characteristics allow birds to respire efficiently but may lead to inhalation of air pollutants at a greater rate than mammals. Thus, in comparison with mammals, birds may be more susceptible to the harmful effects of air pollutants and more likely to be affected at both the population and the community level.

It has been estimated that 48% of all bird species are experiencing population declines (Lees et al., 2022). One of the main drivers of these declines is landcover conversion and degradation associated with urbanization (Tscharntke & Batáry, 2023). With expected increases in urbanization, it is of vital importance to determine how we can achieve bird conservation within urban areas. Although the relationships between urbanization and bird communities have been studied before (Meffert & Dziock, 2013; Melles, 2005; Strohbach et al., 2009; Sultana et al., 2021), few studies have been conducted over longer time scales (> 2 years) or in residential, urban neighborhoods. Moreover, studies assessing the influence of air pollution on bird communities are limited. Specifically, a recent review found 203 papers that considered the influence of air pollution on birds and of these papers only 6.4% (n = 13) assessed effects on bird communities (Barton et al., 2023). The studies that have studied the effects of air pollution on bird communities generally focus on a point source of pollution or used coarse grain pollution indices (Barton et al., 2023), while few, if any, studies have incorporated high-resolution neighborhood measurements of air pollution.

In this study, we examined the relationship between bird communities and features of urban environments, such as neighborhood greenness, landscape features such as roads, other impervious surfaces, and urban parks, and air pollution. We examined the influence of these variables on several metrics of bird communities (relative abundance, species richness, and Hill-Shannon diversity index) in an urban, residential neighborhood at multiple scales. Understanding which factors drive urban bird assemblages within residential areas can help inform and enhance conservation efforts for birds in urban environments.

## Study area

### Methods

We conducted our study in the southwest section of Louisville, Kentucky, USA (centered on 38.205677° N, 85.770016° W), the largest metropolitan area within Kentucky (Figure 1). Louisville is situated in north-central Kentucky along the Ohio River where the state borders Indiana. Our study area was located on the northern edge of the Interior Plateau ecoregion (Type III) and had a sloping elevation that drained towards the Ohio River. While much of the Interior Plateau was historically composed of open savannah woodlands and bluestem prairies, the lower elevation portions of the ecoregion in the Ohio River drainage basin supported oak-hickory forested wetlands. Much of the historic land cover in the Interior Plateau has been converted into cropland, pastureland, and urban areas, such as Louisville. Louisville is dominated by impermeable urban land cover such as roads and buildings with only small, fragmented islands of green space remaining. Our study occurred in a 12 km^2^ neighborhood consisting primarily of residential housing along gridded streets, interspersed with patches of green spaces (e.g., parks) and relatively uniform, impermeable surfaces primarily consisting of streets, driveways, and homes. The study area is intersected by a major highway and contains a horse racing track. It is bordered to the east by a railway and industry, to the north by industry, to the west by mixed non-residential land uses, and to the south by a large park and residential area. Vegetation cover is driven by grass and tree cover in private lawns and public right-of-way areas. Aside from parks and schoolyards, there are few large contiguous green areas. This location is a site of an ongoing large-scale greening study, but all data for our analysis were collected prior to commencement of greening activities (Bhatnagar et al., 2023 [preprint]).

**Figure 1.**
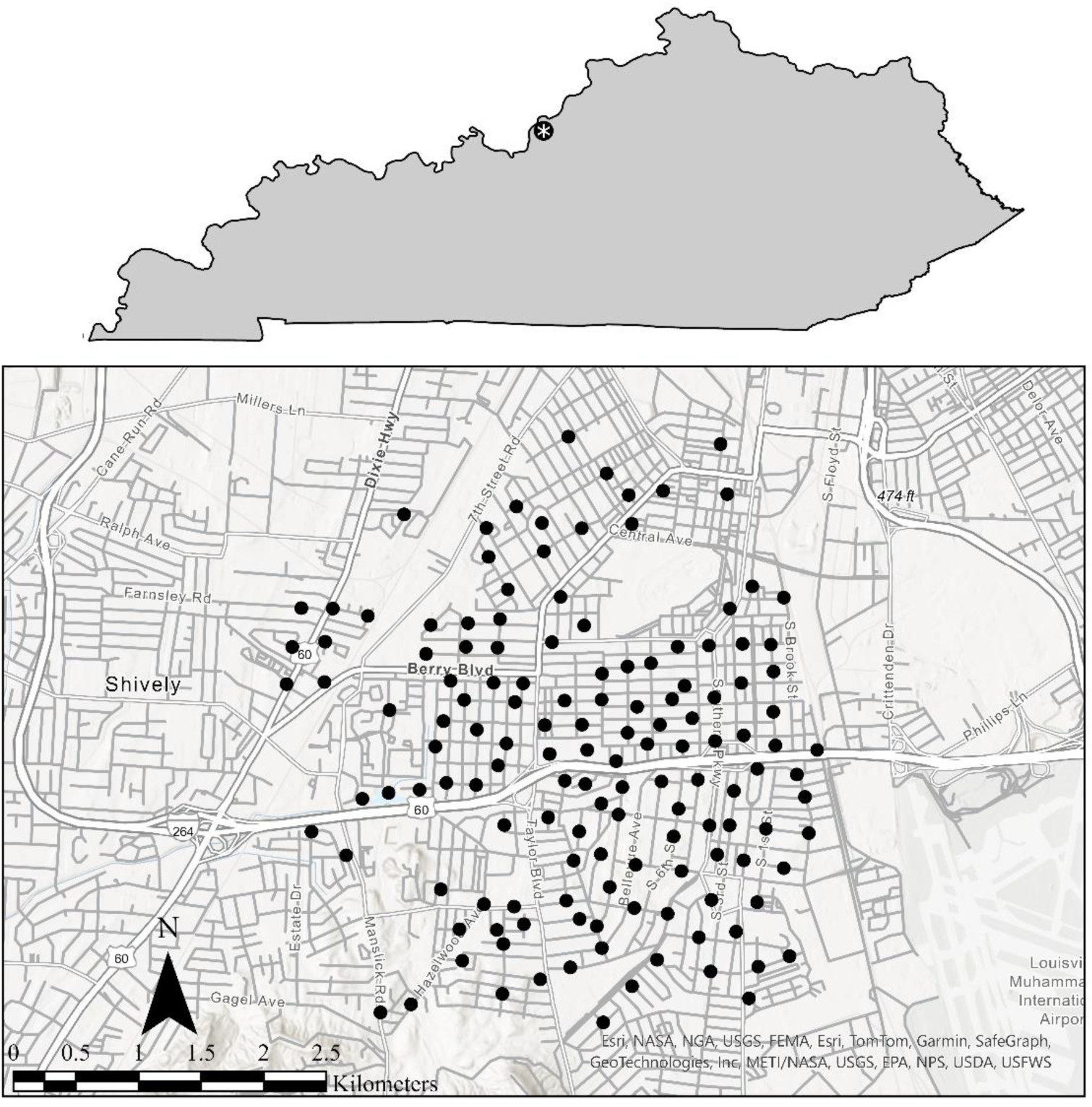
A map of 140 bird point count locations that were sampled as a part of the Green Heart Project from 2019 – 2021 within an urban, residential neighborhood in Louisville, KY, USA.

### Study Design

We established 140 sampling sites within southwest Louisville using systematic random sampling (Figure 1). We designed our study to be able to assess the influence of future greening on wildlife communities. Thus, we established 34 sites within the greening area (i.e., where greening via the planting of trees and other vegetation was planned) and the remaining 106 sites were located approximately every 250 m along concentric rings placed every 250 m from the greening area to 1500 m from the edge to act as control sites. Our stratified random selection of points initially resulted in greater than 140 locations. However, some locations were excluded because of logistical constraints (e.g., safety and accessibility). Following exclusion of those points, we ground-truthed each location and chose a single point with as few obstructions to visibility as possible and within a public right-of-way to act as a stationary point count location for the duration of the study. For sites to be considered independent from one another (i.e., no overlap of surveyed individuals between points), points were spaced 250 m apart according to established standards (Ralph et al., 1995).

### Bird Sampling

To collect bird community data, we surveyed each sample location from 2019 to 2021 between June and August when breeding residents were actively singing. We surveyed each of these sampling points for birds three times within each year with approximately 2 weeks between each survey. To survey points efficiently, the 140 sampling points were assigned to permanent survey groups based on their geographical location. Each group consisted of 10 sampling points that could be reliably sampled throughout one morning, for a total of 14 routes. The points on each of the 14 routes were ordered into the most efficient routes for conducting surveys. Each summer, we randomly selected a starting point for the first visit to each of the 14 survey routes, and then surveyed the remaining points on each route according to our predetermined order. For the subsequent two visits, we rotated the starting point of the route order by three sites each time to reduce the bias of survey timing on results (i.e., surveying the same site at the same time in the day).

We sampled birds at the points located along one route per morning following a standardized point count protocol modified from USGS Breeding Bird Survey methods (Bystrak, 1981). Counts began approximately 5 to 10 min before sunrise and were completed by 4 h past sunrise. Upon arrival at each point, there was a 5-min interval before surveys began to record sampling conditions (e.g., start/end time, start/end temperature, wind, cloud cover, barometric pressure) and to allow birds to acclimate. In the next 10 min, we recorded all birds seen or heard and noted the distance and direction of each bird using a Vortex Ranger 1800 rangefinder and Suunto A-10 compass.

### Bird Community Measures

Though we sampled 140 sites, we excluded 10 sites from the analysis due to greening being initiated at those sites during our sampling period. We combined data from each site across all years and calculated coverage-based estimates of Hill Numbers (q = 0, 1, 2; [Hill, 1973]) with the iNEXT package (Chao et al., 2014) in RStudio version 2022.09.0 (R Core Team, 2021). We evaluated q = 0 or species richness, q = 1 or the exponential of Shannon Diversity Index (hereafter Hill-Shannon diversity), and q = 2 or the inverse Simpson Diversity Index (hereafter Hill-Simpson diversity). As opposed to the original iterations of these diversity indices, Hill number values for Shannon and Simpson indices are expressed in effective species units and greater values are indicative of a more species rich community (Hill, 1973). Calculations of species richness prioritize rarer species, Hill-Simpson diversity prioritizes common species, and Hill-Shannon diversity lies between these two as it does not prioritize rare or common species (Roswell et al., 2021). Our values for Hill-Shannon and Hill-Simpson diversity were highly correlated (Pearson correlation = 0.94). Thus, we only present our analyses of species richness and Hill-Shannon diversity. In addition to these diversity measures, we calculated the average number of invasive and native birds observed across surveys as an index of relative abundance for each site. We did not include species detected beyond 50 m, flyovers, or species likely to have home ranges exceeding the 250 m distance between points (e.g., American Crows [*Corvus brachyrhynchos*] and most birds of prey) in our analysis.

### Environmental Covariates

Our covariates included vegetation, air pollution, and distance-based variables that we selected a priori based on their likely influence on bird communities (Table S1). To determine the most relevant zone of influence, we explored the effects of these variables at multiple scales. We chose to evaluate relationships between our bird community metrics and vegetative and air pollution metrics within buffers with a radius of < 500 m. Specifically, each variable was calculated within 50, 100, 150, 200, 250, or 500 m buffers. The air pollution metrics were not evaluated at 500 m because we did not have air pollution data outside of the study area boundaries. This maximum buffer distance was chosen for two reasons. First, land managers within urban, residential neighborhoods, like our study site, will generally be the most successful in implementation of change at smaller scales. Second, the breeding home ranges of birds we included in our analyses are generally < 250 m and thus we chose to evaluate up to 500 m because it is twice this maximum distance. In addition to these measures, we collected data on the fine-scale vegetation variables within a 50 m buffer around each sampling location and the distance to two landscape attributes that may influence bird communities.

To quantify metrics of greenness, we collected data from the 2018-2019 summer average of cloud-free multispectral images retrieved from Planet Lab’s Planetscope at a 4 m resolution and areal LiDAR at 30 points per m resolution collected from a commissioned fixed-wing areal flyover in 2019. We calculated native-resolution NDVI cover from the multispectral imagery, canopy cover estimates with a machine learning landcover classification algorithm, and Leaf Area Index estimates (LAI) for all canopy areas using multispectral imagery combined with LiDAR point processing and quantification based on the Beer-Lambert law. We then compiled the mean values of NDVI, percent canopy cover, and mean of LAI cover within radial buffer areas by applying the Focal Statistics tool in ESRI ArcGIS Pro, using raster datasets at native resolution as the input to calculate mean buffer values as a raster-based output. To quantify the mean of greenness metrics within buffer areas of sampling points, we extracted the overlaying raster cell values at the corresponding GPS coordinate of each sampling location, which represents the mean value for each metric within a respective buffer area.

Fine-scale vegetation characteristics were measured for all visible vegetation within a 50 m buffer around the bird count point at each of our 140 sites. Within these buffers, all trees were counted and identified to the lowest possible taxonomic classification. The height of all trees was estimated through the tangent method (Larsen et al. 1987) using a Vortex Ranger 1800 rangefinder and Suunto PM-5 clinometer. Additionally, all visible shrubs were counted along a 100 m transect that bisected the buffer through the center point and ran parallel to residential streets. Shrubs were defined as any dense vegetation with obvious understory structure between the canopy and ground layers and were further classified as deciduous or evergreen.

The NO_2_ mixing ratio and ultrafine particle (UFP, airborne particles <100 nm in diameter) number concentration were estimated for each biodiversity sampling location. Oxides of nitrogen passive samplers (Ogawa, Pompano Beach, FL) were deployed for two weeks once every two months at sixty locations across the study area. These samples were extracted and analyzed for NO_x_ (NO + NO_2_) and NO_2_ by ion chromatography. Year 2019 arithmetic mean mixing ratios at each site were used as input to a land use regression model (LUR) to create an NO_2_ concentration surface over the study area. UFP was monitored at 1-second time resolution using a mixing condensation particle counter (Brechtel Model 1720 MCPC, Hayward, CA). The MCPC was deployed on a vehicle and each public road in the study area was driven several times covering various seasons and time of day. The UFP data were post-processed and used as input to an LUR model to create a UFP concentration surface over the study area. Details about the NO_2_ sampling and UFP monitoring, data post-processing, and LUR model implementation have been published (Prathibha, 2021).

The LUR concentration surfaces were used to estimate the NO_2_ and UFP concentrations in a circular buffer around each bird count point. First, LUR concentration surfaces were discretized using a grid with 30 m cells. Next, area-weighted concentrations at each location were calculated for buffers of up to 250 m diameter at 50 m increments. All calculations were performed using a Geographical Information System (ESRI’s ArcGIS Pro 3.0.2, Redlands, CA).

For our spatial characteristics related to the proximity of our points to certain structures or land cover types, shapefile layers were created that identified all green spaces and impermeable surfaces larger than 10 hectares surrounding the study area. Shapefile layers were also created that identified high traffic roads (>5000 vehicles/day) based on the Kentucky Transportation Cabinet database of annual average daily traffic (AADT) from 2019 (KYTC, 2019). For each site, we used the ArcMap distance measuring tools to determine distances (m) between each bird count point and the nearest greenspaces and large impermeable surfaces that were greater than 10 ha and high traffic roads.

### Analyses

We centered and scaled our data prior to analysis to allow for model comparison and assessed spatial autocorrelation in our data using the Moran’s I function in the Program R (R Core Team 2022) package Ape (Paradis and Schliep 2019). Spatial autocorrelation was present for all metrics (Moran’s I: 0.02 – 0.05, P < 0.05), except the average number of invasive birds observed per survey (Moran’s I = 0.003, P = 0.26). Consequently, we ran models to assess the influence of our covariates on each bird community metric using the Program R package SpaMM and the ‘fitme’ function (Rousset & Ferdy, 2014). The ‘fitme’ function accounts for spatial autocorrelation via fitting a Matérn correlation model that includes the X and Y coordinates of each point as a random-slope term. We evaluated model fit of our models with a gaussian, Poisson, negative binomial, and Gamma distribution using the ‘simulateResiduals’ and ‘plot’ functions in the package DHARMa (Hartig, 2022). When appropriate for a specific distribution, each of these models were run with and without a log link. Based on the resulting uniform qq and residual plots, we found that a Gamma distribution with a log link best fit our data and used this distribution for all models.

Initially, we ran separate models to assess relationships between each bird community metric and each independent variable calculated within 50, 100, 150, 200, 250, or, for NDVI, canopy cover, and LAI only, 500 m buffers (Table S1). We also considered the inclusion of each fine-scale and distance-based measure within a separate model set. We included a null model in each model set and compared the model sets using the conditional Akaike’s Information Criteria (cAIC). We incorporated the variable and, when pertinent, given scale within each top model (i.e., with the lowest cAIC) into models considering the independent and additive influence of our covariates on each bird community metric. When the null model was within 2 Δ𝑐𝐴𝐼𝐶 of the top model within each model set, we did not include that variable in the final, additive model.

Additionally, we evaluated correlations between the final variables using the function ggpairs in the Program R package GGally (Schloerke et al., 2021). Any variables with Pearson correlation coefficients > 0.5 were excluded from our final models. Final models within two Δ𝑐𝐴𝐼𝐶 were considered competing models and variables were considered significant predictors if the 95% confidence interval of the beta values did not cross zero.

## Results

Overall, we detected 68 species of birds. After excluding flyovers and species detected beyond 100 m, we observed 43 focal species in the area. Six of the observed species are currently classified as species of greatest conservation need according to the Kentucky Department of Fish and Wildlife Resources (KDFWR) state action plan (KDFWR, 2013). These species were the Green Heron (*Butorides virescens*), Great Egret (*Ardea alba*), Peregrine Falcon (*Falco peregrinus*), American kestrel (*Falco sparverius*), Dickcissel (*Spiza americana*), and Wood Thrush (*Hylocichla mustelina*).

None of the fine-scale or distance-based metrics we evaluated were associated with bird species richness, Hill-Shannon diversity, or the relative abundances of native or invasive birds (Table S2). Species richness (q = 0) was negatively associated with NO_2_ levels at all buffer sizes (Table 1). The relationship between species richness and NO_2_ levels was strongest at the 200 m scale. Conversely, NDVI had a positive relationship with species richness at the 50 m scale only (Table 1). Canopy cover influenced species richness at the 50 and 500 m scales while LAI influenced species richness at 50 and 150 m (Table 1). Our final, combined model included both NO_2_ at 200 m and NDVI at 50 m, indicating that in our study site, both factors are important drivers of species richness (Table 2). Specifically, we found a positive and negative relationship between bird species richness and NDVI and NO_2_, respectively (Tables 3, Figure 2). Like species richness, NO_2_ levels negatively influenced Hill-Shannon diversity (q = 1) at all scales.

**Table 1.**
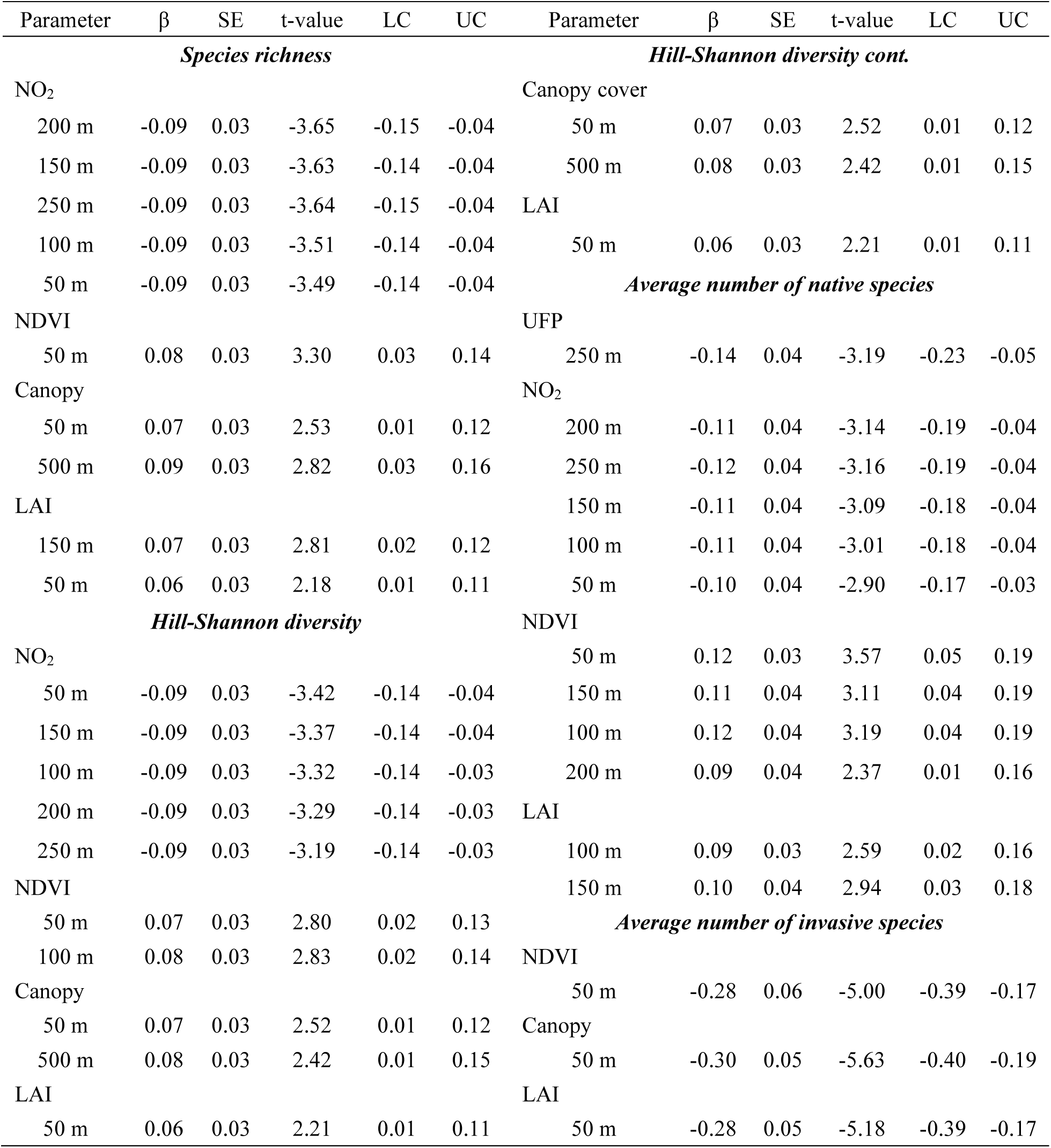
Beta coefficients (β) and associated standard errors (SE), t-values, and lower (LC) and upper (UC) 95% confidence intervals associated with models considering the independent effects of vegetation (normalized difference vegetation index [NDVI], canopy cover, and leaf area index [LAI]) and air pollution (nitrogen dioxide [NO_2_] and ultrafine particulate matter [UFP]) at multiple scales on bird species richness, Hill-Shannon diversity, and the average number of native and invasive bird species in an urban, residential neighborhood in Louisville, KY, USA. Data are not shown if the null model was the top model, the null model was within two delta conditional Akaike’s Information Criteria (ΔcAIC) of the top model, or for any competing models that were within two ΔcAIC of the null model within each model set. The cAIC and ΔcAIC for all models within each set are provided in Supplementary Table 2.

**Table 2.** The conditional Akaike’s Information Criteria (cAIC) and delta cAIC (ΔcAIC) associated with models considering the influence of vegetation (normalized difference vegetation index [NDVI] and canopy cover) and air pollution (nitrogen dioxide [NO_2_] and ultrafine particulate matter [UFP] on bird species richness, Hill-Shannon diversity, and the average number of native and invasive bird species in an urban, residential neighborhood in Louisville, KY, USA.

| Metric | cAIC | ΔcAIC |
| --- | --- | --- |
| *Species richness ($q = 0$)* |  |  |
| **NO<sub>2</sub> (200 m) + NDVI (50 m)** | **443.40** | **0.00** |
| NDVI (50 m) | 446.38 | 2.97 |
| NO<sub>2</sub> (200 m) | 446.77 | 3.36 |
| Null | 453.57 | 10.17 |
| *Hill-Shannon diversity ($q = 1$)* |  |  |
| **Canopy (50 m) + NO<sub>2</sub> (50 m)** | **387.45** | **0.00** |
| NO<sub>2</sub> (50 m) | 390.65 | 3.20 |
| Canopy (50 m) | 394.61 | 7.16 |
| Null | 398.99 | 11.54 |
| *Average number of native species* |  |  |
| **NDVI (50 m) + UFP (250 m) + NO<sub>2</sub> (250 m)** | **399.18** | **0.00** |
| **NDVI (50 m) + UFP (250 m)** | **400.11** | **0.92** |
| NDVI (50 m) + NO<sub>2</sub> (250 m) | 401.52 | 2.33 |
| UFP (250 m) + NO<sub>2</sub> (250 m) | 403.98 | 4.79 |
| NDVI (50 m) | 405.76 | 6.58 |
| UFP (250 m) | 406.12 | 6.94 |
| NO<sub>2</sub> (250 m) | 406.58 | 7.40 |
| Null | 412.77 | 13.58 |
| *Average number of invasive species* |  |  |
| **Canopy (50 m)** | **561.78** | **0.00** |
| Null | 584.89 | 23.11 |

**Figure 2.**
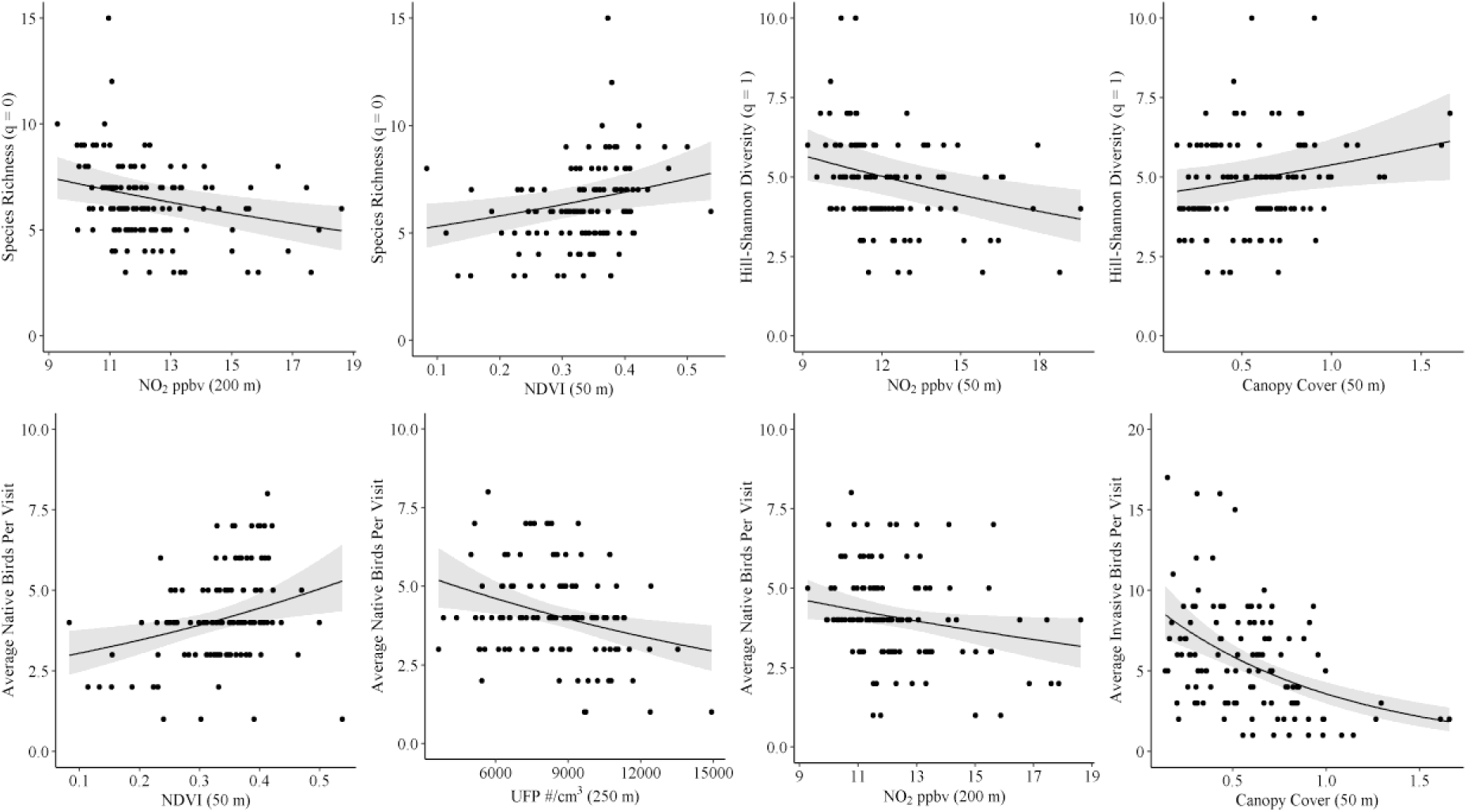
Relationships between vegetation (normalized difference vegetation index [NDVI] or canopy cover) and air pollution (nitrogen dioxide [NO_2_] and ultrafine particulate matter [UFP]) metrics and bird species richness, Hill-Shannon diversity, and the average number of native and invasive bird species in an urban, residential neighborhood in Louisville, KY, USA.

However, the strongest relationship between Hill-Shannon diversity and NO_2_ levels occurred at 50 m. The NDVI and canopy cover correlated with Hill-Shannon diversity at the 50 and 100 m and 50 and 500 m scale, respectively, while LAI only correlated with Hill-Shannon diversity at the 50 m scale (Table 1). Based on our final, combined model, canopy cover and NO_2_ at 50 m correlated positively and negatively, respectively, with Hill-Shannon diversity (Tables 2 and 3, Figure 2).

**Table 3.** Beta coefficients (β) and associated standard errors (SE), t-values, and lower (LC) and upper (UC) 95% confidence intervals associated with models considering the influence of vegetation (normalized difference vegetation index [NDVI] and canopy cover) and air pollution (nitrogen dioxide [NO_2_] and ultrafine particulate matter [UFP]) on bird species richness, Hill-Shannon diversity, and the average number of native and invasive bird species in an urban, residential neighborhood in Louisville, KY, USA.

| Metric | $\beta$ | SE | t-value | LC | UC |
| --- | --- | --- | --- | --- | --- |
| **Species richness ($q = 0$)** |  |  |  |  |  |
| NO<sub>2</sub> (200 m) | -0.08 | 0.03 | -2.99 | -0.13 | -0.03 |
| NDVI (50 m) | 0.07 | 0.03 | 2.57 | 0.01 | 0.12 |
| **Hill-Shannon diversity ($q = 1$)** |  |  |  |  |  |
| NO<sub>2</sub> (50 m) | -0.08 | 0.03 | -3.18 | -0.13 | -0.03 |
| Canopy cover (50 m) | 0.06 | 0.03 | 2.23 | 0.01 | 0.11 |
| **Average number of native species** |  |  |  |  |  |
| *Top model* |  |  |  |  |  |
| NDVI (50 m) | 0.10 | 0.03 | 2.84 | 0.03 | 0.16 |
| UFP (250 m) | -0.10 | 0.04 | -2.78 | -0.18 | -0.03 |
| NO<sub>2</sub> (200 m) | -0.07 | 0.03 | -2.12 | -0.14 | 0.00 |
| *Competing model* |  |  |  |  |  |
| NDVI (50 m) | 0.11 | 0.03 | 3.34 | 0.04 | 0.18 |
| UFP (250 m) | -0.12 | 0.04 | -2.95 | -0.20 | -0.04 |
| **Average number of invasive species** |  |  |  |  |  |
| Canopy cover (50 m) | -0.30 | 0.05 | -5.63 | -0.40 | -0.19 |

The average number of native bird species was associated with LAI, NDVI, NO_2_, and UFPs at multiple scales (Table 1). However, canopy cover did not influence the relative abundance of native birds at any scale (Table S2). Our top model included NDVI at 50 m, UFPs at 250 m, and NO_2_ at 200 m and we had one competing model that only included NDVI at 50 m and UFPs at 250 m (Table 2). We found a negative relationship between UFPs at 250 m and NO_2_ levels at 200 m and the average number of native birds (Tables 2 and 3, Figure 2). Conversely, we observed a positive relationship between the average number of native birds and NDVI at 50 m (Tables 2 and 3, Figure 2). The average number of invasive species was negatively associated with LAI, canopy cover, and NDVI at the 50 m scale only (Table 1). The relationship between canopy cover at 50 m and the relative abundance of invasive birds was strongest (Tables 2 and 3, Figure 2). We found no relationships between invasive bird species and UFPs or NO_2_ levels at any scale (Table S2).

## Discussion

Our findings offer novel insights into the associations between avian communities, vegetation cover, and air pollution in an urban setting. Previous studies have reported relationships between various measures of vegetation complexity (e.g., NDVI, canopy cover, and LAI) and bird diversity and abundances in urban settings (Bar-Massada et al., 2012; Bushra et al., 2021; Debinski et al., 2006; Lerman, 2011; Sandström et al., 2006; Threlfall et al., 2016).

However, our comprehensive spatial assessment found that vegetation metrics are largely associated with bird community measures at relatively fine spatial scales (generally, 50 m) and that the average number of invasive birds observed was only associated with greenness at the 50 m scale. Moreover, unlike the relative abundances of native birds, the relative abundances of invasive birds (i.e., largely House Sparrows [*Passer domesticus*] and European Starlings [*Sturnus vulgaris*]) was negatively associated with all vegetation metrics at 50 m, suggesting that a greener urban setting favors native bird communities. This negative relationship is likely because House Sparrows and European Starlings both prefer open, disturbed areas (Cabe, 2020; Lowther & Cink, 2020).

Our results suggest birds, particularly native species, avoid areas with greater levels of air pollutants such as NO_2_ and UFP. These findings are similar to the results of a study in Tunisia, which found that bird abundances and diversity decrease with increasing proximity to a factory complex, which they considered a point source of pollution (Alaya-Ltifi & Selmi, 2014).

Additionally, the authors reported that House Sparrows became more dominant with increasing proximity to the factory complex. A similar study found bird diversity declined with increasing distance from several stone crusher units, which correlated with changes in the air pollutants that were quantified in our study (Saha & Padhy, 2011). Bird densities and diversity also correlate with heavy metal pollution along a pollution gradient (Belskii & Mikryukov, 2018) and proximity to copper smelter sites (Eeva et al., 2012). Despite these past studies, our fine-scale measures of NO_2_ and UFPs across our study site have allowed us to disentangle and evaluate the effects of air pollution versus other factors (i.e., vegetation) on bird communities.

The differences in bird diversity and relative abundances we observed relative to NO_2_ and UFP may be explained by direct toxic effects of these pollutants. Additionally, birds may detect air pollution and alter their space use (Eeva & Lehikoinen, 1998). Birds also avoid elevated levels of noise associated with traffic (Ortega, 2012), which is the main source of these pollutants on our study site. Given exposure to air pollution could adversely affect the health of the bird, reducing exposure times may be an important strategy to maximize fitness. Such avoidance behavior may also be due to adverse conditions such as smell or acute physiological responses such as difficulties in breathing or stress. For instance, in American Kestrels, exposure to a combination of pollutants, including NO_2_ at 2 ppm, led to an increase in the plasma levels of corticosterone (CORT) and ethoxyresorufin-O-deethylase (EROD) (Cruz-Martinez et al., 2015). Plasma CORT indicates stress, whereas elevated EROD levels usually represent detoxification effort. Similarly, when exposed to a chemical mixture that included NO_2_ at 2 ppm, American Kestrels showed a decrease in their hypothalamus-pituitary-thyroid axis activity, which can influence their metabolism as well as their reproduction (Fernie et al., 2016). Exposure to air pollutants could also induce respiratory stress. For instance, it has been reported that exposure to air pollution associated with urban environments negatively influences the pulmonary function of Rock Doves (*Columba livia*) (Lorz & López, 1997), and that Rock Doves homed faster in more polluted areas, which may be an attempt to decrease exposure to polluted air (Z. Li et al., 2016). However, more research is needed to evaluate the impact of air pollution on bird behavior. Finally, a negative relationship between noise pollution and bird communities has been reported (Ortega, 2012). Given the influence of traffic on the air pollution metrics we measured, it is possible that noise pollution is the underlying driver of the bird community responses that we observed in our study. Additionally, NO_2_ may be a proxy for other traffic-related air pollutants (TRAPs) that could be responsible for the bird community responses we observed.

We conducted our study during the breeding season when the acquisition of arthropods is important to all breeding birds. During this period, arthropods are a main food source for young birds regardless of an adult’s foraging guild (i.e., insectivorous versus granivorous; (O’Connor, 1984). The nestling stage is energy intensive as nestlings are in a phase of rapid growth and therefore reduction in food resources can significantly reduce their growth rate and ultimately affect their future fitness (Grames et al., 2023; O’Connor, 1984). The results of previous studies suggest that arthropods are negatively influenced by air pollution. Specifically, point pollution sources correlate with a decrease in terrestrial arthropod abundances and reduced arthropod body sizes (Zvereva & Kozlov, 2010). Consequently, birds may have less access to arthropods near polluted areas and the arthropods they can feed on are smaller, which would require birds to acquire more food to feed their young. Thus, availability of food resources may be an important driving factor in the patterns we observed between bird community metrics and the local levels of NO_2_ and UFP.

To the best of our knowledge, this is one of the few studies to consider the influence of high-quality greenness measures and fine-scale, site-specific pollution measures on bird communities. However, our study has some limitations. These include a non-representative sample and a lack of noise pollution measures. In the future, it will be valuable to incorporate noise pollution measures to tease apart the relative influences of noise versus air pollution on bird communities. Despite these limitations, our results demonstrate that fine-scale (50 m) manipulation of vegetation, which would be the easiest to accomplish within an urban, residential neighborhood, is key to conservation of bird communities within these urban neighborhoods. Moreover, given the inverse relationships of vegetation metrics with native and invasive bird abundances, greening efforts at finer scales should benefit native birds via increasing structural complexity and potentially reducing competition with invasive birds for food and cover resources. Moreover, our study demonstrates that reductions in NO_2_ at any scale are likely to benefit bird diversity and the relative abundances of native birds. Conversely, efforts to reduce the influence of UFP on native birds should focus at the broadest scale (250 m) we evaluated. Unlike native bird abundances, invasive bird abundances were not sensitive to NO_2_ or UFP. Thus, reductions in air pollution, in combination with greening that increase NDVI, canopy cover, and LAI within an urban neighborhood are likely to increase the abundances of native birds while reducing the abundances of invasive birds.

For a manager in an urban environment, implementation of greening requires obtaining support, and ultimately permission, from multiple private landowners. Given the responses of the bird communities we observed occurring at the finest scales (50 m), this suggests that fine-scale manipulation of vegetation via greening even in small portions of urban, residential neighborhoods are likely to benefit bird communities. Finally, avian communities contribute to the environmental microbiota (Rook, 2013; Teixeira et al., 2013) and increase natural sound profiles, both of which may promote human health (Buxton et al., 2021; Rook, 2013; Teixeira et al., 2013). Therefore, our findings of an association between higher avian biodiversity with greenness may represent one link in the complex pathway through which greenness imparts extensive benefits on the health of nearby residents.

## Acknowledgements

This work was supported in part by grants from the National Institute of Health (ES 029846 and ES023716), The Nature Conservancy, the University of Louisville’s Center for Healthy Air Water and Soil, The Kentucky Ornithological Society, the Kentucky Chapter of The Wildlife Society, and Murray State University’s Jones College of Science Engineering and Technology and Watershed Studies Institute. We are thankful to Yan He for post-processing the air pollution geospatial results, the many field assistants that have been involved in collecting data associated with this research, and to Christina Lee Brown for her in-kind support of the field team.

