## Supplementary Tables for "Drivers of Bird Communities in an Urban Neighborhood Vary by Scale"

Supplementary Table 1. Air pollution and vegetation metrics collected to evaluate factors influencing bird communities in an urban, residential neighborhood in Louisville, KY, USA.

| Metric |
|---|
| *Air pollution* |
| Ultrafine particulate matter (#/cm$^3$) |
| Nitrogen dioxide (ppm) |
| *Greenness* |
| Normalized vegetation index |
| Canopy cover (%) |
| Leaf area index |
| *Shrub density (#/m)* |
| Deciduous |
| Evergreen |
| Combined |
| *Distance (m) from* |
| High traffic roads (>5000 vehicles/day) |
| All parks |
| Parks > 10 ha |
| Impervious surface > 10 ha |
| *Tree height (m)* |
| Average |
| Maximum |

Supplementary Table 2. Conditional Akaike's Information Criteria (cAIC) and ΔcAIC values for models.

| Species Richness (q = 0) | | | Shannon-Hill Diversity (q = 1) | | | Average Number of Native Species | | | Average Number of Invasive Species | | |
|---|---|---|---|---|---|---|---|---|---|---|---|
| Parameter | cAIC | ΔcAIC | Parameter | cAIC | ΔcAIC | Parameter | cAIC | ΔcAIC | Parameter | cAIC | ΔcAIC |
| **UFP** | | | | | | | | | | | |
| UFP (250 m) | 453.38 | 0.00 | UFP (250 m) | 397.81 | 0.00 | UFP (250 m) | 406.12 | 0.00 | Null | 584.89 | 0.00 |
| Null | 453.57 | 0.19 | UFP (200 m) | 398.26 | 0.45 | UFP (200 m) | 408.15 | 2.03 | UFP (100 m) | 586.17 | 1.28 |
| UFP (200 m) | 453.81 | 0.42 | UFP (150 m) | 398.56 | 0.74 | UFP (150 m) | 409.66 | 3.55 | UFP (50 m) | 586.62 | 1.73 |
| UFP (150 m) | 454.56 | 1.18 | UFP (100 m) | 398.90 | 1.08 | UFP (100 m) | 411.27 | 5.15 | UFP (150 m) | 586.84 | 1.94 |
| UFP (50 m) | 454.69 | 1.31 | Null | 398.99 | 1.17 | UFP (50 m) | 411.96 | 5.84 | UFP (200 m) | 587.05 | 2.16 |
| UFP (100 m) | 455.74 | 2.36 | UFP (50 m) | 399.68 | 1.87 | Null | 412.77 | 6.65 | UFP (250 m) | 587.08 | 2.19 |
| **$NO_2$** | | | | | | | | | | | |
| $NO_2$ (200 m) | 446.77 | 0.00 | $NO_2$ (50 m) | 390.65 | 0.00 | $NO_2$ (200 m) | 406.58 | 0.00 | $NO_2$ (250 m) | 583.29 | 0.00 |
| $NO_2$ (150 m) | 446.85 | 0.08 | $NO_2$ (150 m) | 391.19 | 0.54 | $NO_2$ (250 m) | 406.58 | 0.00 | $NO_2$ (200 m) | 583.47 | 0.19 |
| $NO_2$ (100 m) | 447.51 | 0.74 | $NO_2$ (100 m) | 391.40 | 0.75 | $NO_2$ (150 m) | 406.86 | 0.28 | $NO_2$ (150 m) | 583.50 | 0.22 |
| $NO_2$ (50 m) | 447.57 | 0.80 | $NO_2$ (200 m) | 391.69 | 1.04 | $NO_2$ (100 m) | 407.21 | 0.63 | $NO_2$ (100 m) | 583.87 | 0.59 |
| $NO_2$ (250 m) | 446.93 | 0.16 | $NO_2$ (250 m) | 392.34 | 1.68 | $NO_2$ (50 m) | 407.63 | 1.05 | $NO_2$ (50 m) | 584.06 | 0.78 |
| Null | 453.57 | 6.81 | Null | 398.99 | 8.33 | Null | 412.77 | 6.19 | Null | 584.89 | 1.61 |
| **NDVI** | | | | | | | | | | | |
| NDVI (50 m) | 446.38 | 0.00 | NDVI (50 m) | 395.09 | 0.00 | NDVI (50 m) | 405.76 | 0.00 | NDVI (50 m) | 565.44 | 0.00 |
| NDVI (100 m) | 449.47 | 3.09 | NDVI (100 m) | 396.35 | 1.26 | NDVI (150 m) | 405.98 | 0.22 | NDVI (100 m) | 576.03 | 10.58 |
| NDVI (150 m) | 450.06 | 3.68 | NDVI (150 m) | 397.36 | 2.28 | NDVI (100 m) | 406.54 | 0.78 | NDVI (150 m) | 580.16 | 14.72 |
| NDVI (200 m) | 452.90 | 6.52 | NDVI (200 m) | 398.62 | 3.54 | NDVI (200 m) | 409.97 | 4.21 | NDVI (200 m) | 583.30 | 17.86 |
| Null | 453.57 | 7.20 | Null | 398.99 | 3.90 | NDVI (250 m) | 411.47 | 5.71 | Null | 584.89 | 19.45 |
| NDVI (250 m) | 454.74 | 8.37 | NDVI (250 m) | 399.40 | 4.32 | Null | 412.77 | 7.00 | NDVI (250 m) | 584.93 | 19.49 |
| NDVI (500 m) | 456.13 | 9.75 | NDVI (500 m) | 400.80 | 5.72 | NDVI (500 m) | 414.38 | 8.62 | NDVI (500 m) | 585.74 | 20.30 |
| **Canopy** | | | | | | | | | | | |
| Canopy (50 m) | 450.25 | 0.00 | Canopy (50 m) | 394.61 | 0.00 | Canopy (150 m) | 411.02 | 0.00 | Canopy (50 m) | 561.78 | 0.00 |
| Canopy (500 m) | 451.96 | 1.71 | Canopy (500 m) | 396.43 | 1.82 | Null | 412.77 | 1.75 | Canopy (100 m) | 577.82 | 16.04 |
| Canopy (150 m) | 452.56 | 2.31 | Canopy (100 m) | 397.78 | 3.17 | Canopy (100 m) | 413.06 | 2.05 | Canopy (150 m) | 581.26 | 19.48 |
| Canopy (200 m) | 452.81 | 2.57 | Canopy (200 m) | 398.01 | 3.41 | Canopy (200 m) | 413.47 | 2.46 | Canopy (200 m) | 582.00 | 20.22 |
| Canopy (100 m) | 453.29 | 3.04 | Canopy (150 m) | 398.20 | 3.60 | Canopy (50 m) | 413.57 | 2.56 | Canopy (500 m) | 583.67 | 21.89 |
| Null | 453.57 | 3.33 | Canopy (250 m) | 398.45 | 3.84 | Canopy (250 m) | 413.72 | 2.71 | Canopy (250 m) | 584.54 | 22.76 |
| Canopy (250 m) | 453.88 | 3.63 | Null | 398.99 | 4.38 | Canopy (500 m) | 414.34 | 3.32 | Null | 584.89 | 23.11 |

| Species Richness (q = 0) | | | Shannon-Hill Diversity (q = 1) | | | Average Number of Native Species | | | Average Number of Invasive Species | | |
|---|---|---|---|---|---|---|---|---|---|---|---|
| Parameter | cAIC | ΔcAIC | Parameter | cAIC | ΔcAIC | Parameter | cAIC | ΔcAIC | Parameter | cAIC | ΔcAIC |
| **LAI** | | | | | | | | | | | |
| LAI (150 m) | 451.31 | 0.00 | LAI (50 m) | 396.15 | 0.00 | LAI (100 m) | 407.87 | 0.00 | LAI (50 m) | 564.18 | 0.00 |
| LAI (50 m) | 451.47 | 0.16 | LAI (150 m) | 397.15 | 1.00 | LAI (150 m) | 409.78 | 1.91 | LAI (150 m) | 575.01 | 10.83 |
| LAI (200 m) | 452.10 | 0.79 | LAI (100 m) | 397.48 | 1.33 | LAI (200 m) | 412.18 | 4.31 | LAI (100 m) | 577.16 | 12.98 |
| LAI (100 m) | 452.75 | 1.44 | LAI (200 m) | 397.57 | 1.42 | LAI (50 m) | 412.33 | 4.46 | LAI (200 m) | 579.03 | 14.85 |
| LAI (500 m) | 453.19 | 1.88 | LAI (500 m) | 397.89 | 1.74 | Null | 412.77 | 4.89 | LAI (250 m) | 583.38 | 19.20 |
| Null | 453.57 | 2.27 | LAI (250 m) | 398.66 | 2.51 | LAI (250 m) | 413.02 | 5.15 | LAI (500 m) | 583.45 | 19.28 |
| LAI (250 m) | 453.80 | 2.49 | Null | 398.99 | 2.84 | LAI (500 m) | 413.90 | 6.03 | Null | 584.89 | 20.71 |
| **Other measures** | | | | | | | | | | | |
| Shrub | 452.70 | 0.00 | Decid. shrub$^2$ | 397.38 | 0.00 | Shrub$^2$ | 411.28 | 0.00 | Evergr. shrub | 584.20 | 0.00 |
| Decid. shrub$^2$ | 453.44 | 0.74 | Evergr. shrub | 397.38 | 0.00 | Decid. shrub | 412.69 | 1.40 | Avg tree height | 584.36 | 0.17 |
| All parks | 453.55 | 0.85 | Max Tree | 398.19 | 0.81 | Null | 412.77 | 1.48 | Max Tree | 584.39 | 0.19 |
| Null | 453.57 | 0.87 | Shrub | 398.73 | 1.35 | Decid. shrub$^2$ | 413.85 | 2.57 | Evergr. shrub$^2$ | 584.86 | 0.66 |
| Max. tree height | 453.65 | 0.95 | Evergr. shrub$^2$ | 398.75 | 1.37 | Impervious dist. | 413.87 | 2.58 | Null | 584.89 | 0.70 |
| Shrub$^2$ | 454.34 | 1.64 | All parks | 398.92 | 1.54 | Shrub | 413.92 | 2.64 | Decid. shrub$^2$ | 585.21 | 1.01 |
| High traffic road | 454.38 | 1.67 | Null | 398.99 | 1.61 | High traffic road | 414.08 | 2.79 | Shrub$^2$ | 585.30 | 1.10 |
| Decid. shrub | 454.81 | 2.11 | High traffic road | 400.16 | 2.78 | Max. Tree | 414.10 | 2.82 | All parks | 586.05 | 1.85 |
| Evergr. shrub | 455.02 | 2.31 | Shrub$^2$ | 400.37 | 2.99 | Parks > 10 ha | 414.15 | 2.87 | High traffic road | 586.19 | 1.99 |
| Parks > 10 ha | 455.06 | 2.35 | Avg. tree height | 400.41 | 3.03 | Evergr. shrub | 414.23 | 2.95 | Impervious dist. | 586.36 | 2.16 |
| Avg. tree height | 455.53 | 2.82 | Parks > 10 ha | 400.49 | 3.12 | All parks | 414.25 | 2.97 | Parks > 10 ha | 586.40 | 2.20 |
| Impervious dist. | 455.92 | 3.22 | Decid. shrub | 400.89 | 3.52 | Evergr. shrub$^2$ | 414.31 | 3.02 | Shrub | 586.41 | 2.21 |
| Evergr. shrub$^2$ | 456.43 | 3.72 | Impervious dist. | 401.00 | 3.63 | Avg. tree height | 414.55 | 3.27 | Decid. shrub | 586.56 | 2.36 |
